# Interpreting patterns of X chromosomal relative to autosomal diversity in aye-ayes (*Daubentonia madagascariensis*)

**DOI:** 10.1101/2025.01.25.634876

**Authors:** John W. Terbot, Vivak Soni, Cyril J. Versoza, Mark Milhaven, Adriana Calahorra-Oliart, Devangana Shah, Susanne P. Pfeifer, Jeffrey D. Jensen

## Abstract

We here present high-quality, population-level sequencing data from the X chromosome of the highly-endangered aye-aye, *Daubentonia madagascariensis*. Using both polymorphism- and divergence-based inference approaches, we quantify fine-scale mutation and recombination rate maps, study the demographic and selective processes additionally shaping variation on the X chromosome, and compare these estimates to those recently inferred from the autosomes in this species. Results suggest that an equal sex ratio is most consistent with observed patterns of variation, and that no sex-specific demographic patterns are needed to fit the empirical site frequency spectrum. Further, reduced rates of recombination were observed relative to the autosomes as would be expected, whereas mutation rates were inferred to be similar. Utilizing the estimated population history together with the mutation and recombination rate maps, we evaluated evidence for both recent and recurrent selective sweeps as well as balancing selection across the X chromosome, finding no significant evidence supporting the action of these episodic processes. Overall, these analyses provide new insights into the evolution of the X chromosome in this species, which represents one of the earliest splits in the primate clade.

## Introduction

Different chromosomes within a genome are generally considered to be effectively unlinked (Morgan 1911), such that each may be treated as an independent sample of the evolutionary history of the population in question. Following from this, autosomes may be utilized as pseudo-replicates when performing population genomic analyses (e.g., demographic estimation). However, due to their association with sex determination and sex-associated differences, sex chromosomes often require separate consideration. The extent and impact of these effects will naturally differ based on the form of sex determination, the age of the sex chromosome system, the population sex ratio, and numerous other contributing factors (for a discussion, see the reviews of Ellegren 2011; Bachtrog *et al*. 2014).

Mammals are generally characterized by an XY sex-determination system in which the heterogametic sex (XY) develops into males and the homogametic sex (XX) develops into females. Outside of a relatively small pseudoautosomal region (PAR), no recombination occurs between the X and Y chromosomes (unless otherwise stated, discussion of the “X chromosome” in this study is restricted to the larger, non-pseudoautosomal region (non-PAR) most impacted by the unique properties of sex chromosomes). Additionally, oogenesis is typically restricted to a single developmental period, while spermatogenesis occurs throughout the lifespan of an organism. These factors alone could be expected to lead to differences in evolutionarily-important factors including rates of mutation and recombination, particularly as compared to autosomes in which chromosomal copy number is identical between the sexes. For example, there are fewer copies of the X chromosome relative to an autosome (3/4 under even sex ratios), generally leading to a lower effective population size, resulting in differing degrees of genetic drift and modifying demographic scaling on the X (e.g., Pool and Nielsen 2007; Singh *et al*. 2007). Relatedly, sex-specific migration may further modify demographic patterns on the X relative to the autosomes (Laporte and Charlesworth 2002). Secondly, sex chromosomes uniquely experience a prolonged portion of their history in a haploid state, meaning that when in males, mutations will be directly visible to selection even whilst rare and regardless of dominance (Haldane 1924; Charlesworth *et al*. 1987; and see Bachtrog *et al*. 2009). Moreover, Hartl (1971,1972) noted that the change in mean fitness at X-linked loci experiencing equivalent selection in both sexes would be greater than for autosomal loci. Thirdly, owing to differences in the number of germ cell divisions in males and females, male-biased mutation patterns have been widely observed (e.g., Haldane 1946; Hurst and Ellegren 1998; McVean 2000).

In primates specifically, numerous insights have been gained with regards to X chromosome evolution, often discussed relative to the expected 3:4 ratio of effective population sizes on the X:autosomes. Notably, if there exists a higher male variance in reproductive success, this ratio may exceed the expected 0.75, reaching as high as 1.125 when there is effectively a single reproducing male; if the variance in female reproductive success is higher, the ratio may approach 0.5625 when there is effectively a single reproducing female (Charlesworth 2001). As might be expected, the genetic diversity on the sex chromosomes of humans have been most heavily studied (see the review of Webster and Wilson Sayres 2016), with results indicating, for example, deviation in this ratio of autosomal to X-linked genetic diversity relative to the 0.75 expectations (e.g., Keinan *et al*. 2009), as well as likely differences between the historical effective population sizes of males and females (e.g., Wilder *et al*. 2004). Outside of humans, the X:autosome ratio of diversity in primates has been observed to be particularly low in gorillas and orangutans (Prado-Martinez *et al*. 2013), but nearer to a neutral, equal sex ratio expectation in Tonkean macaques (Evans *et al*. 2014).

As the great majority of previous studies have been focused upon the great apes, or biomedically-relevant species, large swathes of the primate clade have remained poorly studied in this regard. Bringing these analyses to a strepsirrhine, we here consider the evolutionary history of the X chromosome of the aye-aye (*Daubentonia madagascariensis*), a nocturnal lemur endemic to Madagascar (Ancrenaz *et al*. 1994; Randimbiharinirina *et al*. 2019). While they are a behaviorally and morphologically unique species, the most relevant features to this study are those pertaining to their sexual characteristics. Morphologically, there is little difference between male and female aye-ayes outside of primary sex characteristics like gonads (Sterling 1993), and in terms of diet and typical behavior, males and females are also largely alike (Ancrenaz *et al*. 1994). There is little evidence of biased sex ratios at birth or during juvenile development, aside from a perhaps subtle bias towards male offspring from younger sires (Tanaka *et al*. 2019). However, there are notable differences in the territorial ranges associated with each sex. While both males and females are known to have sizable home ranges (Randimbiharinirina *et al*. 2019), those of males are thought to be larger and more likely to overlap with the ranges of multiple females and even other males (Sterling 1993; Ancrenaz *et al*. 1994). If this for example were to result in fewer reproductively contributing males, this distinction could be expected to reduce male relative to female effective population sizes.

In order to investigate the evolutionary dynamics of the X chromosome in this sexually monomorphic species, we have here obtained high-coverage sequence data of X chromosomes from five individuals (three females and two males). Usefully, there have been considerable advances in our understanding of the evolutionary genomics of aye-ayes over the past year, which have here served to greatly improve our resolution of evolutionary processes acting on the X. These developments have included the generation of an annotated chromosome-level aye-aye genome (Versoza and Pfeifer 2024), the recent estimation of a demographic history for this population based on the autosomal site frequency spectrum (SFS; Terbot *et al*. 2025), genomic scans for evidence of positive and balancing selection across the autosomes (Soni *et al*. 2024), as well as high-quality pedigree-based estimates of autosomal mutation (Versoza *et al*. 2025) and recombination (Versoza, Lloret-Villas *et al*. 2024) rates, and structural variant architecture (Versoza *et al*. 2024). More recently, indirect estimates of autosomal mutation and recombination rates have been made, utilizing divergence data and patterns of linkage disequilibrium (Soni, Versoza *et al*. 2024), respectively, and the shape of the distribution of fitness effects (DFE) characterizing exonic divergence across the autosomes has additionally been inferred (Soni *et al*. 2025). These studies all provide a novel and exceptional framework in which to perform comparative analyses of the evolutionary dynamics acting on the autosomes relative to the, as of yet unexplored, X chromosome in this species.

Thus, we here present analyses investigating the demographic history of the aye-aye X chromosome relative to the autosomal genome, utilize this demographic history to quantify likely sex ratios in this species, employ forward-in-time simulations to account for the DFE and the resulting background selection effects across the X when model-fitting, estimate fine-scale mutation and recombination rate maps across the chromosome, and perform both polymorphism- and divergence-based scans for functional regions evolving under recurrent positive or balancing selection. Taken together, this study thus represents the first comprehensive investigation of X chromosome dynamics in this highly endangered primate.

### Estimating the demographic history of the X chromosome

Our initial step was to determine if the demographic history previously estimated from the autosomes (Terbot *et al*. 2025) remained consistent with the X chromosome, or whether a history of sex-specific population dynamics would be necessary to explain observed patterns of variation. To this end, following the approach outlined in Terbot *et al*. (2025), we first mapped sequencing reads from five individuals (three females and two males) to the species-specific reference genome (Versoza and Pfeifer 2024) using BWA-MEM v.0.7.17 (Li and Durbin 2009) and called single nucleotide polymorphisms (SNPs) using the Genome Analysis Toolkit (GATK) Best Practices workflow (van der Auwera and O’Connor 2020). To determine the location of the completely sex-linked segment of the X chromosome – the focus of this study – we used differences in male:female coverage to identify the pseudoautosomal boundary separating the non-PAR from the PAR. Notably, earlier work was unable to reliably identify the size of the PAR in aye-ayes (Shearn *et al*. 2020), owing to the noisiness of medium coverage data from a single representative of each sex mapped to the highly-fragmented genome of the distantly-related gray mouse lemur (*Microcebus murinus*, Mmur_2.0 consisting of >10,000 scaffolds). However, the recently released, highly-contiguous genome assembly for the species allowed us to confidently identify this region (Figure 1a).

**Figure 1:**
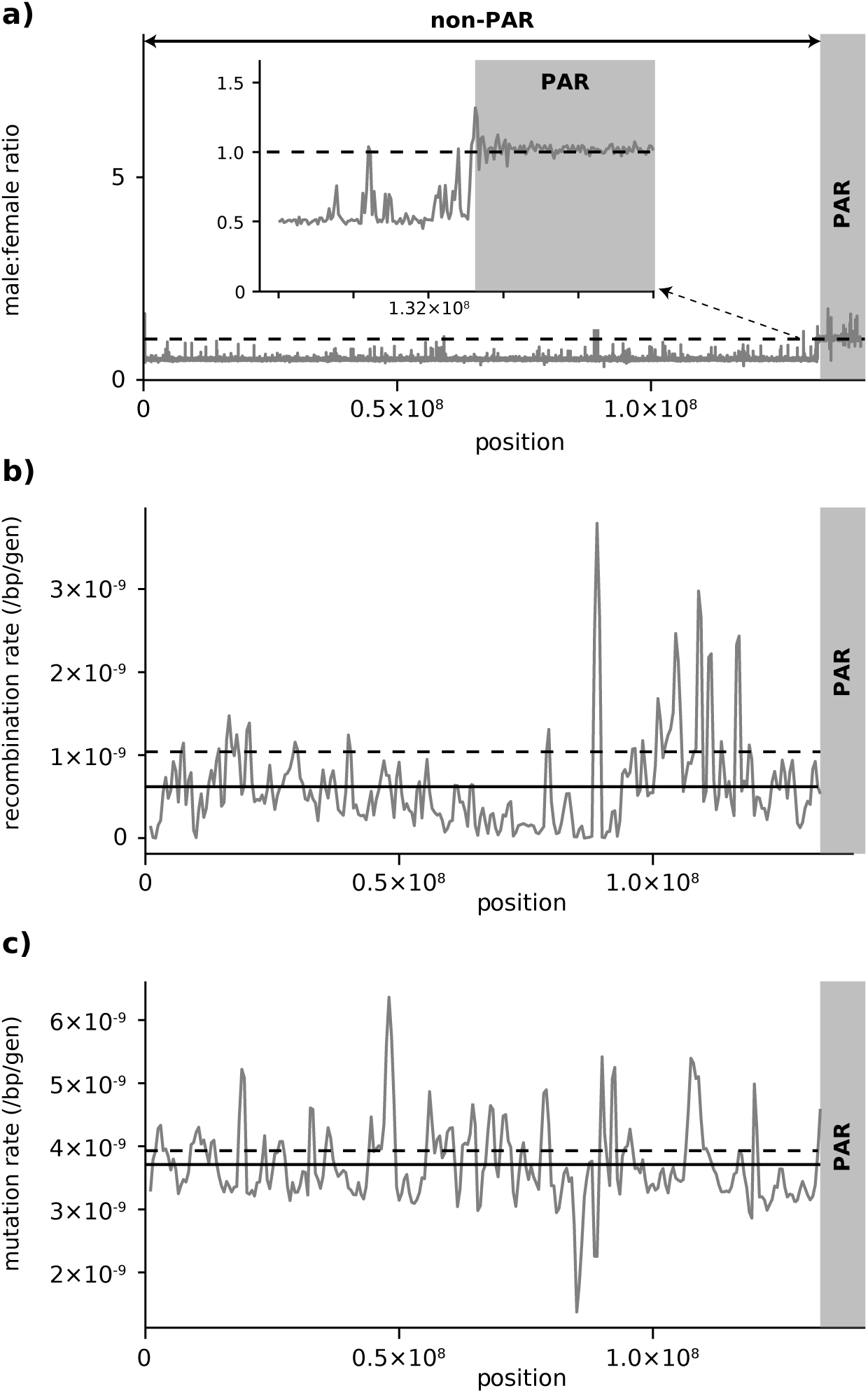
(a) Male:female sequence coverage ratios on the X chromosome were used to identify the pseudoautosomal boundary, separating the pseudoautosomal region (PAR) from the completely sex-linked non-PAR. In the former, a male:female coverage ratio of 1 (indicated by a dashed line) is expected as males and females carry the same number of copies of that region. (b) Recombination and (c) mutation rate maps across the non-PAR. The solid line on each panel indicates the average rates for the X chromosome; the dashed line on each panel indicates the genome-wide average rates for the autosomal chromosomes (Soni, Versoza *et al*. 2024).

As all X chromosomal samples were collected from the “non-North” deme described in Terbot *et al*. (2025), we anticipated that our sample should similarly appear unstructured, which was re-confirmed using ADMIXTURE v1.3 (Alexander *et al*. 2009) (data not shown). We thus used SLiM v.4.3 (Haller and Messer 2023) to simulate 100 X chromosomes under the previously estimated autosomal demographic history for the non-North deme, utilizing a DFE for exonic regions previously estimated in humans as a reasonable proxy of expected purifying and background selection effects (Johri *et al*. 2023; Soni and Jensen 2025). As prior work found the mutation rate in males to be ∼2.7 times greater than that of females (Versoza *et al*. 2025), along with the difference in time spent in males relative to females for the X chromosome, we calculated an average rescaled rate of 2.22 x 10^-8^ mutations per site per generation for males, and 0.82 x 10^-8^ mutations per site per generation for females. Mutations were modeled for the entire non-PAR of the X chromosome as being strictly neutral in non-exonic regions, and drawn from a DFE in exonic regions. From these simulations, variant call files (in .vcf format) of five samples (three females and two males) were produced to mimic the empirical data. Repetitive and constrained regions of these files were then masked to match empirical masking, and SFS were produced. When compared to the empirically observed SFS (Figure 2), the autosomal demographic model scaled for the X was found to match well, suggesting that an alternative X chromosome-specific demographic history is not needed to fit the empirical data.

**Figure 2:**
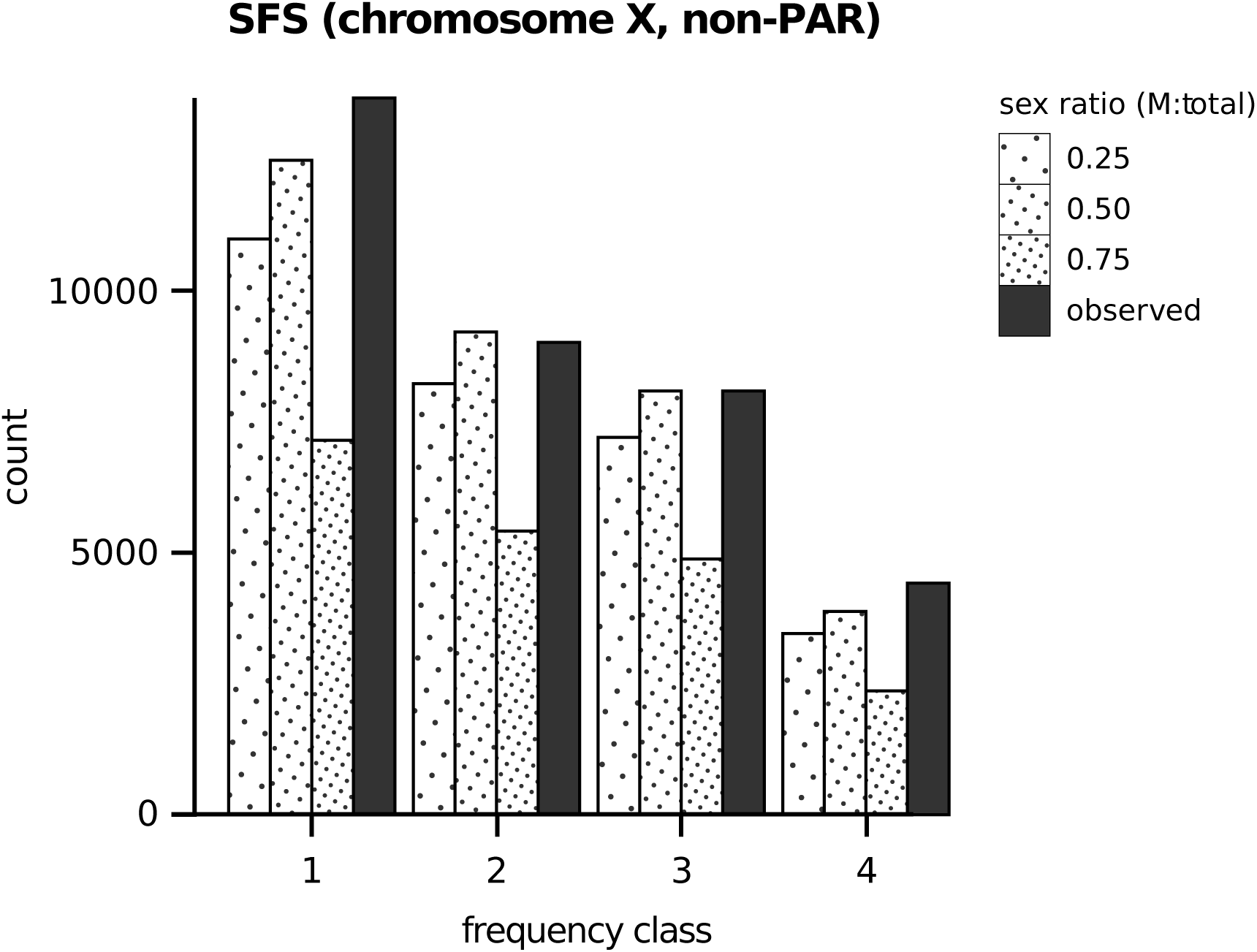
Site frequency spectra (SFS) of neutral sites in the non-pseudoautosomal region (non-PAR) of the X chromosome for the observed data (filled), and average values for simulated data at different sex ratios (0.25 male (M) vs male+female (total), sparsely dotted; 0.5 M:total, moderately dotted; 0.75 M:total, heavily dotted).

Utilizing this demographic history, we next inferred the likely sex ratios characterizing the long-term evolutionary dynamics of this population. As shown (Figure 2), a 1:1 male:female ratio was found to be most consistent with the data, with alternative sex ratios generally reducing variation below the empirical observation, and skewing the SFS away from that observed. These relative differences between alternative sex ratios are consistent with expectations as have been described via simulation (e.g., Spurley and Payseur 2025).

### Inferring a fine-scale recombination rate map of the X chromosome

Following the approach outlined in Soni, Versoza *et al*. (2024), we applied several stringent filter criteria prior to estimating recombination rates across the X chromosome. Specifically, to match the data of the autosomal chromosomes, we removed variants that exhibited an excess of clustering (≥3 variants within 10bp) or heterozygosity (Hardy-Weinberg equilibrium *p*-value < 0.01), or that were located within regions blacklisted by ENCODE (Dunham *et al*. 2012). Afterward, population-scale recombination rates (*ρ*) were inferred across the X chromosome using the LDhat v.2.2 program (McVean *et al*. 2004; Auton and McVean 2007) in interval mode, following the protocols for non-human primates (Auton, Fledel-Alon, Pfeifer, Venn *et al*. 2012; Stevison *et al*. 2016; Pfeifer 2020; Soni, Versoza *et al*. 2024). As shown in Figure 1b, the population-scaled recombination rate for the X chromosome was found to be 0.04 *ρ*/kb, approximately 70% lower than the rate previously reported for autosomes (0.15 *ρ*/kb; Soni, Versoza *et al*. 2024). This reduction is in agreement with the notion that sex chromosomes have suppressed recombination relative to autosomes, owing partly to the lower effective population size and absence of recombination whilst in males (Charlesworth 2017; Olito and Abbott 2023).

### Inferring a fine-scale mutation rate map of the X chromosome

Following the approach of Soni, Versoza *et al*. (2024), we replaced the previously existing low-coverage aye-aye X chromosome from the Kuderna *et al*. (2024) 447-way mammalian alignment with the high-quality, annotated aye-aye X chromosome of Versoza and Pfeifer (2024) using the Cactus pipeline (Armstrong *et al*. 2020). Using this updated multiple-species alignment, substitutions along the aye-aye branch were extracted in both neutral and functional regions, removing any divergent sites observed to be polymorphic. Neutral divergence rates were calculated by considering the number of divergent sites relative to accessible sites in a given window, and exonic divergence by considering the number of divergent sites relative to exonic length. Rates were calculated in windows of 1kb, 10kb, 100kb, and 1Mb, with step sizes equal to half the window size. In order to convert neutral divergence into estimated per-generation mutation rates, a divergence time of 54.9 million years (Horvath *et al*. 2008; Soni *et al*. 2025) was divided by a generation time of 5 years (Ross 2003; Louis *et al*. 2020). As shown, the neutral mutation rate inferred across the X chromosome is consistent with indirectly inferred autosomal rates (Figure 1c), suggesting a mean rate of 3.78 x 10^-9^ mutations per base pair per generation.

### Scans for positive and balancing selection on the X chromosome

In order to investigate the potential of recent, rare bouts of positive or balancing selection, we utilized the demographic model described above to determine statistical thresholds for performing genomic scans. To evaluate evidence for recent selective sweeps, we utilized SweepFinder2 v1.0 (Nielsen *et al*. 2005; DeGiorgio *et al*. 2016) on each SNP in the 100 simulated X chromosomes (-*su*). From this, we established the maximum Composite Likelihood Ratio (CLR) for the aye-aye X chromosome under the estimated population history, and accounting for purifying and background selection effects in coding regions (resulting in a maximum observed value of 217.60). The maximum empirical CLR observed (130.11) was considerably below this threshold (Figure 3a), suggesting that, if recent sweeps have occurred on the X chromosome, they are not identifiable within the context of this evolutionary baseline model (see Poh *et al*. 2014; Johri *et al*. 2022).

**Figure 3:**
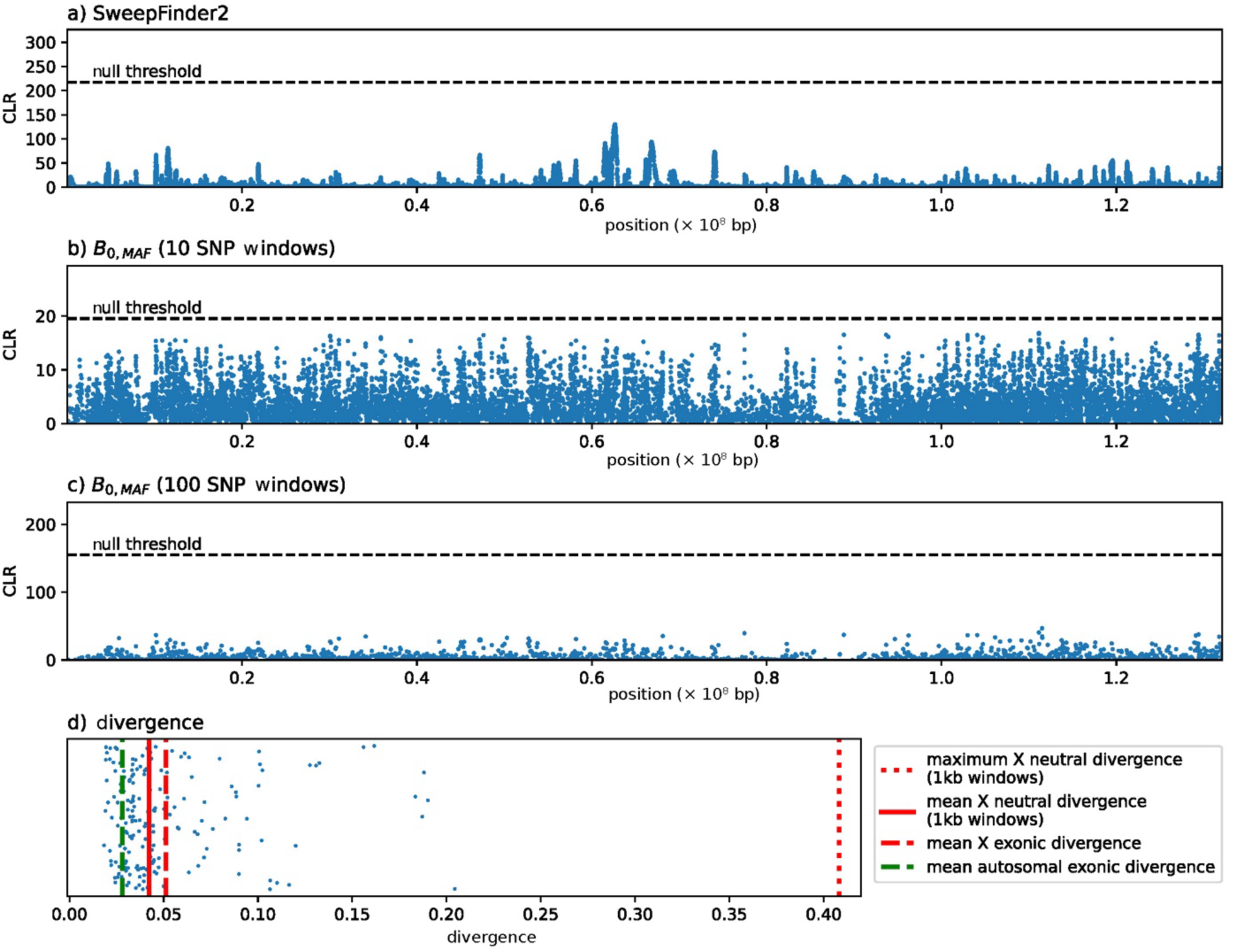
Polymorphism- and divergence-based scans for selection on the X chromosome using (a) SweepFinder2, (b), BalLerMix in 10 SNP windows, and (c) BalLerMix in 100 SNP windows, in which null thresholds (dashed lines) for composite likelihood ratios (CLR) were determined via simulation. (d) Divergence across the X chromosome, with each point representing an X-linked exon. The mean exonic divergence on the X (dashed red line), the mean and maximum neutral divergence on the X in 1kb windows (solid and dotted red lines, respectively), and the mean autosomal exonic divergence (dashed green line) are provided for comparison.

We similarly evaluated evidence for recent balancing selection, utilizing BalLerMix v.2.5 (Cheng and DeGiorgio 2020) on all simulated X chromosomes using the *B_0,MAF_* method – which uses the folded SFS – with 10 and 100 SNP windows and step sizes of 5 and 50, respectively (*-w 10*,*--step 5* and *-w 100*,*--step 50*). The maximum CLR for the simulated data was 19.54 for 10 SNP windows and 154.98 for 100 SNP windows. By comparison, the maximum CLR values for the observed data was 16.86 for the 10 SNP windows and 47.12 for the 100 SNP windows (Figures 3b and 3c). Thus, the maximum baseline model CLR values were well above the maximum observed CLR values, similarly suggesting a lack of support for the action of balancing selection on the X chromosome. This is in contrast to recent autosomal analyses, which found strong support for long-term balancing selection acting on olfactory-related genes in this species (Soni *et al*. 2024).

Finally, with regards to recurrent and long-term adaptive processes, no exons were observed to be diverging more rapidly than the most rapid neutral observation (Figure 3d), once accounting for windows of like size (Table 1). Taken together with the polymorphism-based scans, these results thus suggest limited evidence for either strong and recent, or rapidly recurrent, adaptive evolution across X-linked genes in the aye-aye genome.

**Table 1:** Mean, standard deviation (std.), and maximum (max) neutral divergence on chromosome X based on window size, for windows containing at least 200 accessible sites. The equivalent values from the autosomes are provided for comparison (Soni, Versoza *et al*. 2024).

| window size | autosomal divergence |  |  | X divergence |  |  |
| --- | --- | --- | --- | --- | --- | --- |
|  | mean | std. | max | mean | std. | max |
| 1kb | 0.0437 | 0.0263 | 0.4941 | 0.0424 | 0.0288 | 0.4082 |
| 10kb | 0.0439 | 0.0165 | 0.2681 | 0.0412 | 0.0235 | 0.3364 |
| 100kb | 0.0432 | 0.0088 | 0.1352 | 0.0415 | 0.0121 | 0.1415 |
| 1Mb | 0.0432 | 0.0059 | 0.0730 | 0.0418 | 0.0080 | 0.0734 |

## Summary

By quantifying fine-scale mutation and recombination rate maps across the X chromosome of the highly-endangered aye-aye, we found mutation rates consistent with those inferred on the autosomes via both direct- and indirect-estimation approaches, and recombination rates greatly reduced relative to the autosomes owing to both the lower effective population size as well as the expected absence of recombination in the non-PAR whilst in males. Furthermore, we found that the demographic history of the X chromosome matches that of the autosomes and thus does not require sex-specific population dynamics, and that an equal male-female ratio is most consistent with the observed data, despite previously described differences in territorial ranges between the sexes. Utilizing this well-fit evolutionary baseline model, we additionally constructed thresholds for performing scans of selection using both patterns of polymorphism and divergence, finding extensive evidence for purifying selection as expected, but no significant evidence in support of positive or balancing selection along the X chromosome. Taken together, these results provide unique insights into the X chromosome dynamics of a strepsirrhine, a quantification of male:female sex ratios otherwise extremely difficult to obtain in this elusive, nocturnal, endangered species, and further support for generally reduced mutation rates in lemurs relative to other primates.

## Acknowledgements

We would like to thank the Duke Lemur Center for providing the aye-aye samples used in this study, and members of the Jensen Lab and Pfeifer Lab for helpful discussion. Computations were performed on the Sol supercomputer at Arizona State University (Jennewein *et al*. 2023) and on the Open Science Grid (Pordes *et al*. 2007; Sfiligoi *et al*. 2009), which is supported by the National Science Foundation and the U.S. Department of Energy’s Office of Science. This is Duke Lemur Center publication # XXXX.

